# GPX3 supports ovarian cancer tumor progression *in vivo* and promotes expression of GDF15

**DOI:** 10.1101/2024.01.24.577037

**Authors:** Caroline Chang, Ya-Yun Cheng, Shriya Kamlapurkar, Sierra R. White, Priscilla W. Tang, Amal T. Elhaw, Zaineb Javed, Katherine M. Aird, Karthikeyan Mythreye, Rébécca Phaëton, Nadine Hempel

## Abstract

**Objective:** We previously reported that high expression of the extracellular glutathione peroxidase GPX3 is associated with poor patient outcome in ovarian serous adenocarcinomas, and that GPX3 protects ovarian cancer cells from oxidative stress in culture. Here we tested if GPX3 is necessary for tumor establishment *in vivo* and to identify novel downstream mediators of GPX3’s pro-tumorigenic function.

**Methods:** GPX3 was knocked-down in ID8 ovarian cancer cells by shRNA to test the role of GPX3 in tumor establishment using a syngeneic IP xenograft model. RNA sequencing analysis was carried out in OVCAR3 cells following shRNA-mediated GPX3 knock-down to identify GPX3-dependent gene expression signatures.

**Results:** GPX3 knock-down abrogated clonogenicity and intraperitoneal tumor development *in vivo*, and the effects were dependent on the level of GPX3 knock-down. RNA sequencing showed that loss of GPX3 leads to decreased gene expression patterns related to pro-tumorigenic signaling pathways. Validation studies identified GDF15 as strongly dependent on GPX3. GDF15, a member of the TGF-β growth factor family, has known oncogenic and immune modulatory activities. Similarly, GPX3 expression positively correlated with pro-tumor immune cell signatures, including regulatory T-cell and macrophage infiltration, and displayed significant correlation with PD-L1 expression.

**Conclusions:** We show for the first time that tumor produced GPX3 is necessary for ovarian cancer growth *in vivo* and that it regulates expression of GDF15. The immune profile associated with GPX3 expression in serous ovarian tumors suggests that GPX3 may be an alternate marker of ovarian tumors susceptible to immune check-point inhibitors.

## Introduction

Ovarian cancer remains the most deadly gynecological malignancy and the fifth-leading cause of cancer-related deaths in women in the United States (1). The five-year survival rate for patients with advanced stage epithelial ovarian cancer remains below 30%, with the development of chemoresistance and high rates of recurrence contributing to these poor outcomes. Poly (adenosine diphosphate-ribose) polymerase (PARP) inhibitors have been added to standard of care based on the homologous recombination deficiency status of high grade serous ovarian cancers, with encouraging results, yet development of resistance to newer therapies, and the lack of response to immunotherapy are continuing challenges associated with treatment.

Cancer cells upregulate their antioxidant enzyme expression to adapt to oxidative stress associated with metastatic progression and changing tumor microenvironments (2, 3). Cells are equipped with a sophisticated antioxidant enzyme system to ensure maintenance of cellular redox homeostasis, which includes the glutathione peroxidase family members. While assessing if expression of antioxidant enzymes in serous adenocarcinomas from TCGA is associated with poor patient outcome, we found that high expression of Glutathione peroxidase 3 (GPX3) is most strongly associated with negative patient outcome among all antioxidant enzymes examined (4). GPX3 is an extracellular selenocysteine-containing protein with antioxidant activity, catalyzing the reduction of hydroperoxides, including hydrogen peroxide (H_2_O_2_) and soluble lipid hydroperoxides in the presence of glutathione. Systemically, GPX3 is primarily produced by the kidneys and is the predominant glutathione peroxidase in plasma. Initially, studies reported that many cancer patients display decreased plasma glutathione peroxidase activity and levels, which is associated with selenium deficiency, suggesting that decreases in plasma GPX3 activity are negatively associated with patient outcome, including ovarian cancer patients (5, 6). However, more recent studies have started to demonstrate that tumor-produced GPX3 is associated with chemoresistance and tumor progression in several cancer types (reviewed in (6)). We previously demonstrated that TCGA serous adenocarcinoma cases fall into distinct groups of high and low GPX3 expression, and that high expression of GPX3 in ovarian cancer specimens is associated with poor patient outcome (4). Other studies have similarly found that GPX3 expression is predictive of poor outcome and associated with chemoresistance (7, 8). We demonstrated that GPX3 is advantageous to tumor cell clonogenicity and anchorage-independent survival, and that GPX3 protects ovarian cancer cells from exogenous sources of oxidative stress, including co-culture in patient derived ascites fluid (4). However, it remains unknown if tumor produced GPX3 is necessary for *in vivo* tumor growth and if GPX3 expression contributes to changes in the tumor transcriptome. In the present work we set out to determine the role of GPX3 on *in vivo* tumor progression using a syngeneic xenograft model and to further study its function in ovarian cancer using an unbiased RNA sequencing approach, which identified GDF15 as a novel GPX3 target gene in ovarian cancer.

## Materials & Methods

### Cell culture and generation of GPX3 knock-down cell lines

Luciferase-expressing mouse ovarian cancer ID8-Luc2 cells were maintained in high glucose Dulbecco’s Modification of Eagle’s Medium (DMEM; Corning, 10-017-CV), 4% fetal bovine serum (FBS; Avantor Seradigm,1500-500), 5 mg/mL insulin (MilliporeSigma, I0516), 5 mg/mL transferrin (MilliporeSigma, T8158), 5 ng/mL sodium selenite (MilliporeSigma, S5261) and 100 units/mL penicillin-streptomycin solution (Gibco, 15140122). Human ovarian cancer NIH:OVCAR3 (OVCAR3) cells were purchased from the American Type Culture Collection (ATCC HTB-161; Manassas, VA) and maintained in RPMI 1640 medium (Gibco, 11875) supplemented with 10% FBS and 0.01 mg/ml bovine insulin (MilliporeSigma, I0516). OVCAR4 (MilliporeSigma) were cultured in RPMI 1640 medium (Corning,10-040-CV) supplemented with 10% FBS, 0.25 U/mL insulin and 2mM glutamine. OVCA433 cells (kindly provided by Dr. Susan K. Murphy, Duke University), were cultured in RPMI 1640 medium (Corning,10-040-CV) supplemented with 10% FBS. All cell lines were maintained in 5% carbon dioxide (CO_2_) at 37°C. Cells are routinely tested for mycoplasma (Captivate Bio, 20-700-20) and authenticated using STR sequencing.

To generate stable GPX3 knock-down cells, lentivirus was packaged in 293FT cells using the ViraPower Kit (Invitrogen, Carlsbad, CA), according to manufacturer’s instructions. Two different short hairpin (shRNA, from Sigma-Aldrich shRNA library; Table 1), scrambled shRNA control (Addgene plasmid #1864), or an empty backbone pLKO.1 control (Addgene plasmid #8453) were used to generate virus and infect OVCAR3 cells or ID8 cells, respectively (4).

**Table 1.**
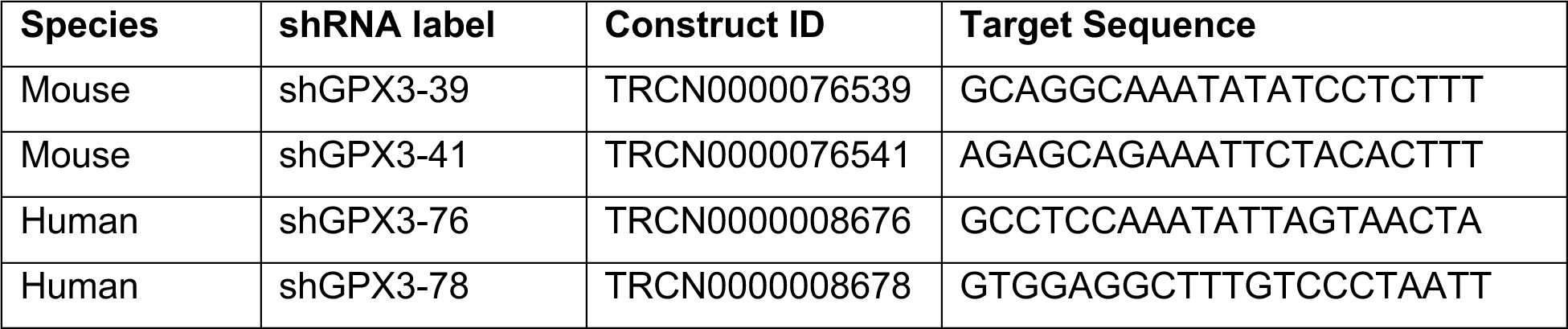
shRNA sequences.

For transient siRNA-mediated GPX3 knock-down cells were seeded to 6 cm dishes at 60-80% confluency. The next day, the non-targeting control (Dharmacon, #D-001810-10-05) or human GPX3 siRNA (Dharmacon, #L-006485-00-0005) were delivered to the cells using Lipofectamine RNAiMAX (Invitrogen, 13778150). After 24 hours, the transfected cells were transferred to 10 cm dishes and incubated for 72 hours. The cells were harvested for RNA isolation and semi-quantitative real-time RT-PCR.

### *In vivo* syngeneic model of ovarian cancer

*In vivo* experiments were approved by the Penn State College of Medicine’s Institutional Animal Care and Use Committee and conformed to the Guide for the Care and Use of Laboratory Animals. Mice were housed in ventilated cages on corncob bedding (7092 Harlan Teklad, Madison, WI), and had *ad lib* access to irradiated rodent chow (2918 Harlan Teklad, Madison, WI) and autoclaved water. The facility was programmed with a 12:12-h light:dark cycle. Room temperature was maintained at 20 ± 2°C, with air humidity range 30 to 70%. Environmental enrichment was provided with an Enviropak (Lab Supply, Fort Worth, Tx). 12 to 16 week old C57BL/6J mice were obtained from Jackson Laboratory and randomly assigned into 3 groups to be injected intraperitoneally (IP) with 1×10^6^ ID8 cells stably expressing either pLKO.1 (n=12), or shRNAs targeting GPX3 shGPX3-39 (n=6), or shGPX3-41 (n=8) resuspended in 0.2 mL sterile phosphate-buffered saline (PBS). Results represent cumulative data from 2 experiments (Exp1: Control n=6, shGPX3-39 n=6, shGPX3-41 n=3; Exp2: Control n=6, shGPX3-41 n=5). Mice were imaged using an IVIS Lumina III In Vivo Imaging System (IVIS; PerkinElmer, Waltham, MA). Mice were injected IP with 10 µL/g of body weight of 15 mg/mL *in vivo* grade D-Luciferin (luciferin; PerkinElmer, Waltham, MA), anesthetized with 2-3% isoflurane in an induction chamber, and then moved to the IVIS imaging chamber and maintained under 2-3% isoflurane for imaging 10 minutes later. Luciferin was prepared and sterilized using a 0.22 µm syringe filter and frozen in 1 mL aliquots until use. Mice were weighed and imaged 1-2 times a week until reaching endpoint. Humane endpoints included significant ascites that affected breathing or movement, severe lethargy, or weight loss greater than 20% of original body weight. Mice were euthanized by CO_2_ asphyxiation followed by cervical dislocation. After euthanasia, abdominal ascites was immediately collected via a sterile 18g needle syringe, and the volumes measured. The omentum was removed, weighed, and imaged using the IVIS in 2 mL PBS and 150 µg/mL luciferin. Tumor samples of the omentum were placed in 600 µL TRIzol Reagent (Invitrogen,15596018) and frozen at −80°C for RNA analysis.

### RNA isolation and semi-quantitative real-time RT-PCR

Total RNA was extracted using the Qiagen RNeasy Mini Kit (Qiagen, Hilden, Germany) or Direct-zol RNA Miniprep kit (Zymo Research, R2052). First strand synthesis was performed using the iScript cDNA Synthesis Kit (Bio-Rad, Hercules, CA) or qScript cDNA Synthesis Kit (Quantabio, 95047), according to manufacturer’s instructions. For mouse tissues, total RNA was isolated from omental tumors after tissues were collected and homogenized in TRIzol Reagent (Invitrogen,15596018). Semi-quantitative real-time RT-PCR was carried out using the Applied Biosystems 7500 Real Time PCR or Bio-Rad CFX Opus 96 Real-Time PCR System with the primer sequences listed below (Table 2), with PowerUp SYBR Green Master Mix (Applied Biosystems) or iTaq Universal SYBR Green Supermix (Bio-Rad). Quantification of each gene was calculated using the ΔΔCt method with values normalized to housekeeping genes (Table 2) and change in transcript expression expressed relative to control cells. For mRNA analysis from tumors, mRNA levels were expressed relative to expression of one control tumor (mouse 1) to demonstrate heterogeneity in expression between tumor specimens.

**Table 2.**
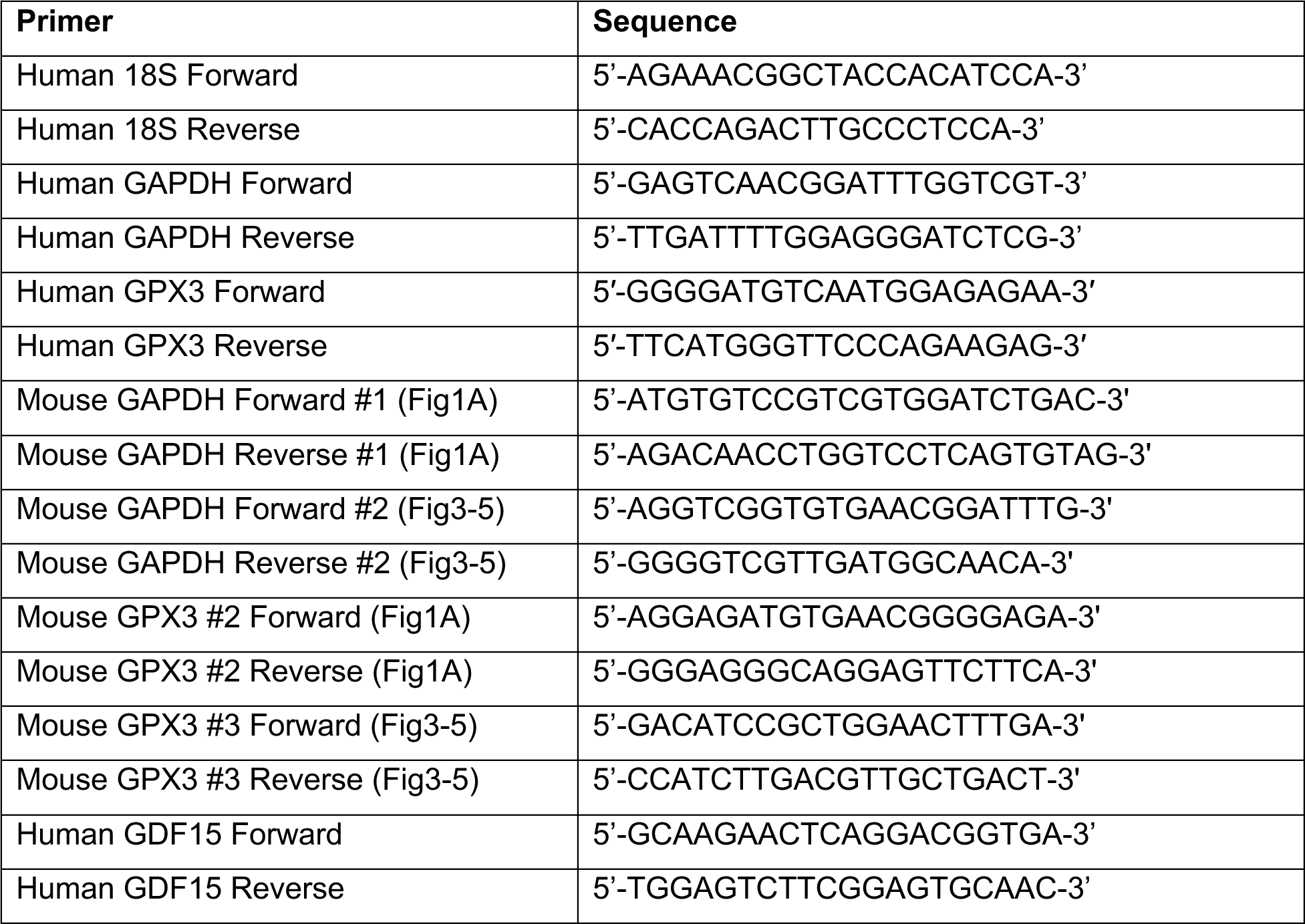
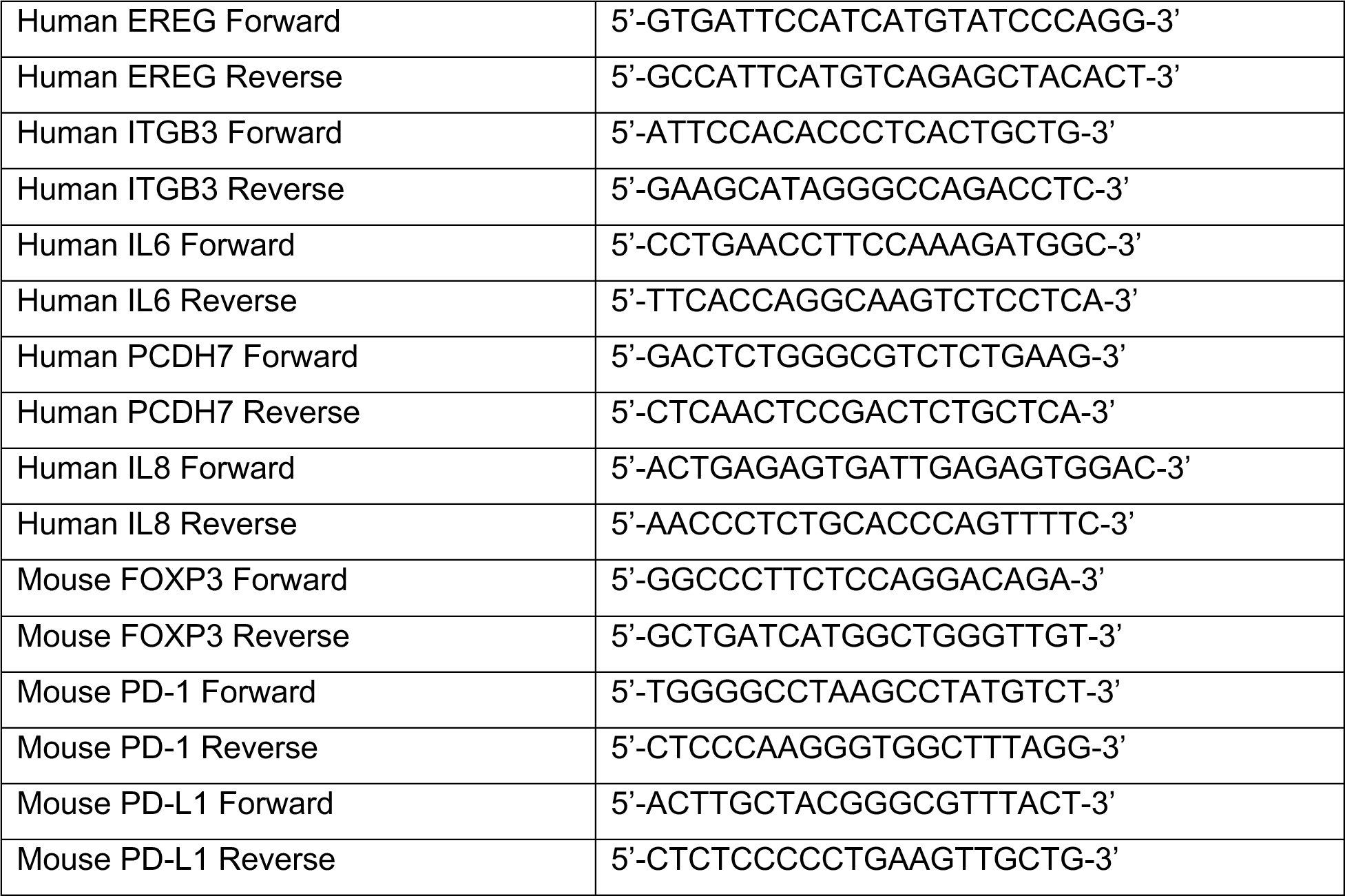
Primer sequences used for semi-quantitative real-time RT-PCR.

### RNA-sequencing

OVCAR3 cells stably expressing pLKO.1 scrambled control, shGPX3-76, or shGPX3-78 were seeded and incubated for 72 hrs. Total RNA was extracted from cells using Direct-zol RNA Miniprep kit (Zymo Research, R2052). mRNA was purified from total RNA using poly-T oligo-attached magnetic beads and first strand cDNA was synthesized using random hexamer primers, followed by second strand cDNA synthesis using dTTP, end repair, A-tailing, adapter ligation, size selection, amplification, and purification for non-directional library preparation (Novogene). RNA sequencing carried out on Illumina NovaSeq 6000 (Novogene) and reads in fastq format processed through perl scripts (Novogene) to obtain clean reads and downstream analyses based on these. Index of the reference genome was built, and paired-end clean reads were aligned to the reference genome hg38 using Hisat2 v2.0.5. FeatureCounts v1.5.0-p3 was used to count the reads numbers mapped to each gene. Then FPKM of each gene was calculated based on the length of the gene and reads count mapped to this gene. Differential expression analysis between control and GPX3 knock-down samples was performed using the DESeq2R package (1.20.0; Supplemental Data Table 1). The resulting P-values were adjusted using the Benjamini and Hochberg’s approach for controlling the false discovery rate (FDR), and genes with an adjusted P-value <0.05 assigned as differentially expressed. Volcano plots were generated using VolcaNoseR software. Gene Set Enrichment Analysis (GSEA) was carried out on presorted DEGs (broadinstitute.org/gsea). Transcription factor analysis and small molecule query L1000CDS2 analysis (LINCS project) was performed using BioJupies (9). RNA sequencing data are accessible through NCBI GEO (GSE254035).

### Clonogenicity assays

ID8 control and GPX3 knock-down cells were seeded in 12 well plates (100 cells per well) and cultured for 10 days under fully supplemented media. After 10 days, 0.05% crystal violet solution (MilliporeSigma, 229288) was used to stain colonies. Colonies were counted using Image J and data expressed as cellular survival fractions.

### Immunoblotting

Cells were harvested by scraping into RIPA buffer (Thermo Scientific, 89901) containing protease and phosphatase inhibitors (Thermo Scientific, 78443). Samples were sonicated (Qsonica) for a total of 5 mins and spun down for 10 mins at 13000 rpm, 4°C. The concentration of the protein samples was measured using Bradford protein assay (Bio-Rad, 5000006). 50 µg of whole cell lysate was used for SDS-PAGE and then transferred to PVDF membranes (Thermo Scientific, PI88520). The membranes were blocked with 5% non-fat milk (Bio-Rad,1706404) in TBS containing 0.1% Tween20 (MilliporeSigma, 900-64-5) for 30 mins, and incubated with primary antibodies at 4°C, overnight (Table 3). Ater washing with TBST (10 mins, 3 times), the membranes were probed with horseradish peroxidase (HRP)-conjugated secondary antibodies at room temperature for 1 hour, and then developed using SuperSignal™ West Femto Maximum Sensitivity Substrate Femto (Thermo Scientifi, 34096). The signals were captured by ChemiDoc XRS+ Imaging System (Bio-Rad).

**Table 3.** Antibodies used.

| **Antibody** | **Manufacturer** | **Cat #** | **Dilution** |
| --- | --- | --- | --- |
| GPX3 | R&D Systems | AF4199 | 1:400 |
| GDF15 | R&D Systems | AF957 | 1:400 |
| Vinculin | Sigma-Aldrich | Aldrich V9131 | 1:1000 |
| Amersham ECL HRP<br>conjugated rabbit IgG | Cytiva | NA934 | 1:10000 |
| Amersham ECL HRP<br>conjugated mouse IgG | Cytiva | NA931 | 1:10000 |
| Mouse anti-goat IgG-HRP | Santa Cruz Biotechnology | sc-2354 | 1:5000 |

For analysis of extracellular GPX3 and GDF15 cells were seeded the previous day and media changed to serum-free and phenol red-free media (Gibco, 11835). Media was collected after 72 hrs, cell debris removed by centrifugation and media concentrated using Amicon Ultra Centrifugal Filters (MilliporeSigma, UFC901024), spinning for 2 hrs at 4000 rpm, 4°C. Protein concentration of the supernatants were quantified by Bradford protein assay (Bio-Rad, 5000006). 30 µg of protein samples were used for immunoblotting with nitrocellulose membranes (Amersham Protran, 10600001) and equal loading demonstrated by Ponceau S staining (MilliporeSigma, P7170-1L).

### Statistical analysis

Data are reported as mean ± SEM from at least three independent experiments. All statistical analyses were chosen based on experimental design, as indicated, and performed using GraphPad Prism 9 (GraphPad Software, San Diego, CA). *p* values < 0.05 were considered significant.

## Results

### GPX3 is important for *in vivo* peritoneal tumor spread in a syngeneic ovarian cancer model

In previous work we demonstrated that GPX3 expression in TCGA ovarian serous adenocarcinoma specimens is associated with poor patient outcome (4). We showed that GPX3 is needed to protect ovarian cancer cells from exogenous oxidative stress. Moreover, GPX3 knock-down decreases clonogenicity when cells are cultured in patient derived ascites fluid and abrogates anchorage-independent survival of OVCAR3 cancer cells (4). To expand on this work, we tested the importance of GPX3 on *in vivo* tumor progression using a syngeneic ovarian cancer xenograft model. Two independent shRNAs targeting GPX3 transcripts were expressed in ID8 mouse ovarian cancer cells and knock-down confirmed by semiquantitative real-time RT-PCR (Figure 1A). Similar to our previous findings in human OVCAR3 cells, GPX3 knockdown in ID8 cells also inhibited their clonogenicity (Figure 1B) (4). The effect on colony formation was dependent on the level of GPX3 knock-down, requiring at least 40% decrease in GPX3 mRNA expression to achieve significant effect on clonogenic survival (Figure 1B).

**Figure 1.**
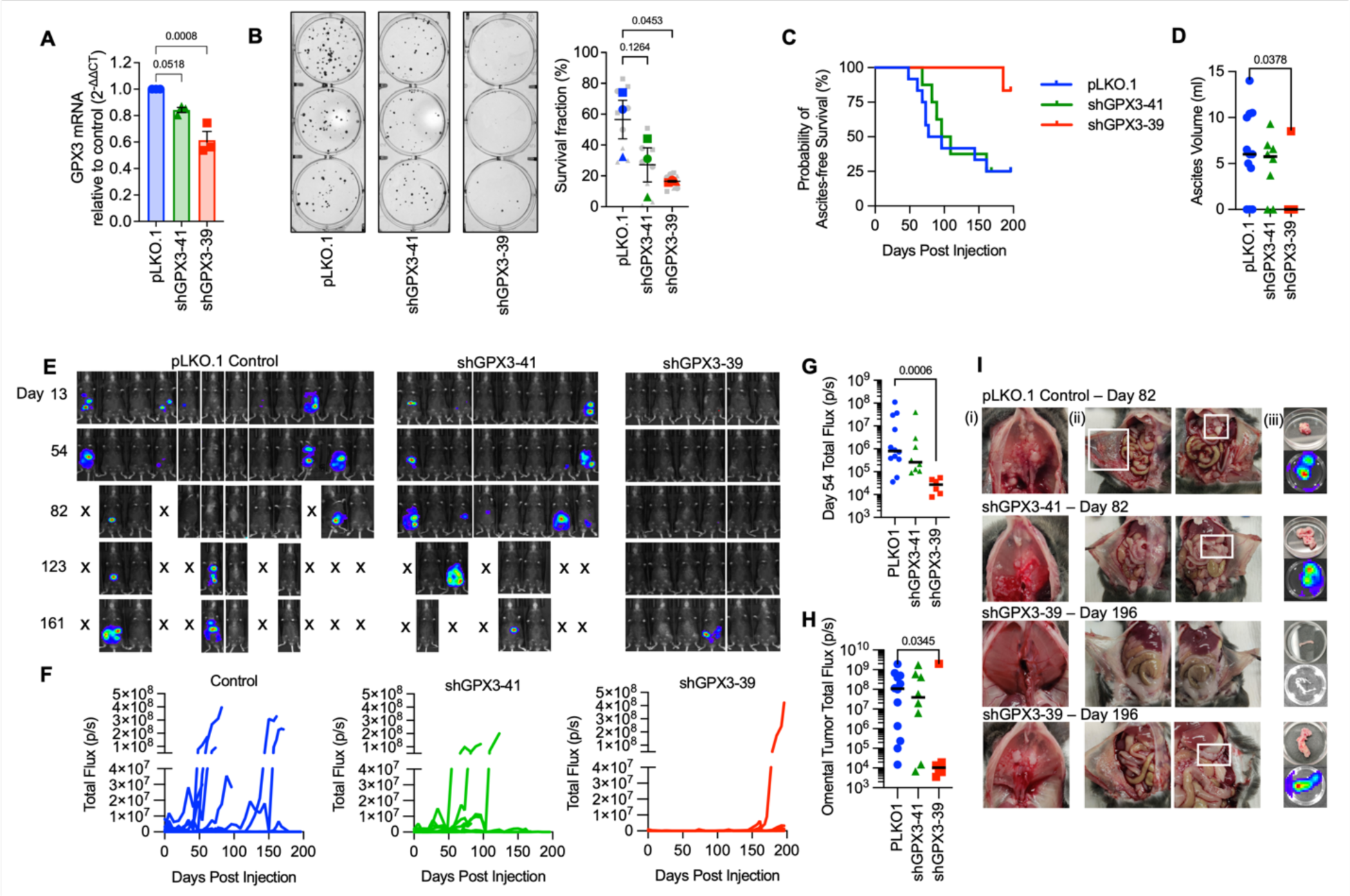
Knock-down of GPX3 decreases tumor burden in a syngeneic mouse ovarian cancer tumor model. A. GPX3 was knocked down using 2 different shRNAs in mouse ID8 tumor cells. GPX3 expression assessed using sqRT-PCR (n=3 One-way ANOVA p=0.0013; Dunnett’s post-test). B. GPX3 decreases clonogenicity of ID8 cells (n=3 biological replicates. Superplot shows underlying technical replicates, One-way ANOVA p=0.0607; Dunnett’s post-test). C. ID8 tumor cells transduced with pLK0.1 control, shGPX3-41 or shGPX3-39 shRNAs were IP injected into immunocompetent C57BL/6J mice. GPX3 knock-down by shRNA-39 results in significant delay in the onset of ascites, which was used as endpoint for survival studies. Kaplan Meier curves show probability of ascites-free survival (pLK0.1 control n=12, shGPX3-41 n= 8, shGPX3-39 n=6; Log-rank Mantel-Cox test, P value = 0.0443). D. Quantification of total ascites volume collected at necropsy (Kruskal-Wallis test). E. Intraperitoneal tumor growth of ID8 tumor cells transfected with pLK0.1 control, shGPX3-41 or shGPX3-39 shRNAs in C57BL/6J mice was monitored using luminescence imaging F. Quantification of individual mouse tumor luminescence signals over time (pLK0.1 control n=12, shGPX3-41 n= 8, shGPX3-39 n=6; Total Flux p/s). G. Median tumor burden as assessed by luminescence on Day 54 (pLK0.1 control n=12, shGPX3-41 n= 8, shGPX3-39 n=6; Total Flux p/s; Kruskal-Wallis test). H. Tumor luminescence of isolated omenta at necropsy (pLK0.1 control n=12, shGPX3-41 n= 8, shGPX3-39 n=6; Kruskal-Wallis test). I. Tumor burden was detected on the diaphragm (i), peritoneal wall (ii) and omentum (iii). Representative images of mice at necropsy endpoint day 82 from pLK0.1control and shGPX3-41 groups are shown. A representative abdomen from the 5/6 mice that did not develop tumors, and the one of 6 mice in the shGPX3-39 group that developed tumors (lower panel) are shown from endpoint day 196.

C57BL/6J mice injected IP with control ID8 cells (pLKO.1) developed omental tumors and abdominal ascites faster than mice injected with either knockdown cell line (Figure 1C-F). Ascites burden was used as an endpoint determinant and median ascites free survival for mice injected with shGPX3-41 ID8 cells was 102.5 days compared to 85.6 days in control tumor cell injected mice (Figure 1E), with 9/12 control mice and 6/8 shGPX3-41 mice developing omental tumors and abdominal ascites (Figure 1D-I). Tumors were also detected on the diaphragm, peritoneal walls, gastrointestinal tract, and the right kidney (Supplemental Figure 1). As seen in clonogenicity studies a dose-dependent effect of GPX3 knock-down was also observed *in vivo*. Injection of ID8 shGPX3-39 cells that display a more significant GPX3 knock-down also resulted in significantly reduced tumor burden compared to the control group, with only 1/6 mice developing ascites (Figure 1D)_and abdominal tumors at endpoint of the study (day 196, Figure 1E-I). The one mouse in the shGPX3-39 group that developed tumors displayed similar-sized omental tumors and ascites burden compared to the control group (Figure 1H, I). The above data suggest that tumor expressed GPX3 is necessary for optimal tumor dissemination *in vivo*.

### Transcriptional changes in response to GPX3 knock-down

To determine how GPX3 knock-down might be aiding tumor progression and affecting intrinsic gene expression changes, we carried out RNA sequencing on human OVCAR3 cells where GPX3 expression was stably knocked down using two independent shRNAs (Figure 2A, Figure 3A). Gene set enrichment analysis (GSEA) analysis revealed that GPX3 expression is associated with a number of cancer-associated gene expression signatures, in particular metabolic and biosynthesis genes, and their associated regulatory signaling pathways, including mTOR, PPAR and AMPK signaling (Figure 2B; Supplemental Figure 2; Supplemental Data Table 1). None of the upregulated gene expression pathways assessed by KEGG or Reactome analysis had an FDR <0.05 in response to knock-down. In agreement, small molecule gene expression signature analysis demonstrated that GPX3 knock-down most mimics the effects PI3K/mTOR inhibitors on gene expression (Figure 2C, Supplemental Data Table 1). Analysis of validated Transcription Factor binding data (ChEA, ENCODE) revealed that the gene signatures down regulated following GPX3 knock-down are targets of CEBPB (immune response, metabolism), ATF3 (stress response), SOX2 (stemness) and FOXA2 (metabolism; Figure 2D; Supplemental Data Table 1). These data suggest that GPX3 expression regulates pro-tumorigenic pathways involved in stress response and metabolism.

**Figure 2.**
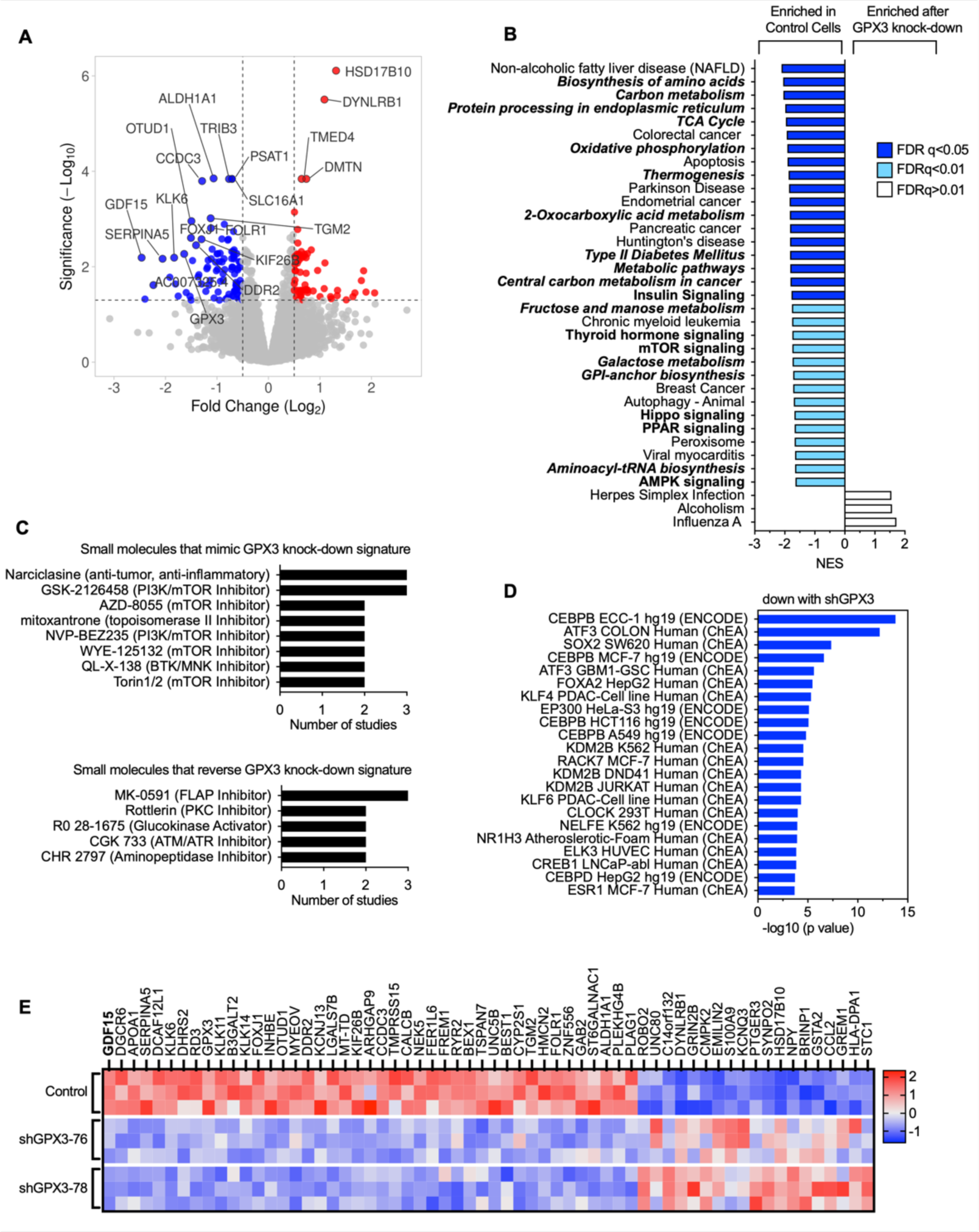
RNA sequencing demonstrates downregulation of pro-tumorigenic signaling pathways in response to GPX3 knock-down in OVCAR3 cells. A. Volcano plot of differentially expressed genes (DEGs) following GPX3 knock-down using two different shRNAs (shRNA-76 & shRNA-78; n=3) relative to scramble shRNA control transduced OVCAR3 cells. B. Gene set enrichment analysis of RNA sequencing data following GPX3 knock-down (KEGG) demonstrates enrichment of tumor associated metabolic pathways in control cells relative to GPX3-knock-down cells. C. Small Molecule Query (L1000CDS^2^) was carried out to identify gene expression signatures that mimic or reverse gene expression changes following GPX3 knock-down. D. Transcription factor enrichment analysis (ENCODE & ChEA) on RNAsequencing data following GPX3 knock-down. E. Heat map of top differentially expressed protein coding genes (DEG) following GPX3 knock-down (z-scores; log 2-fold change >1, p adjust <0.05).

**Figure 3.**
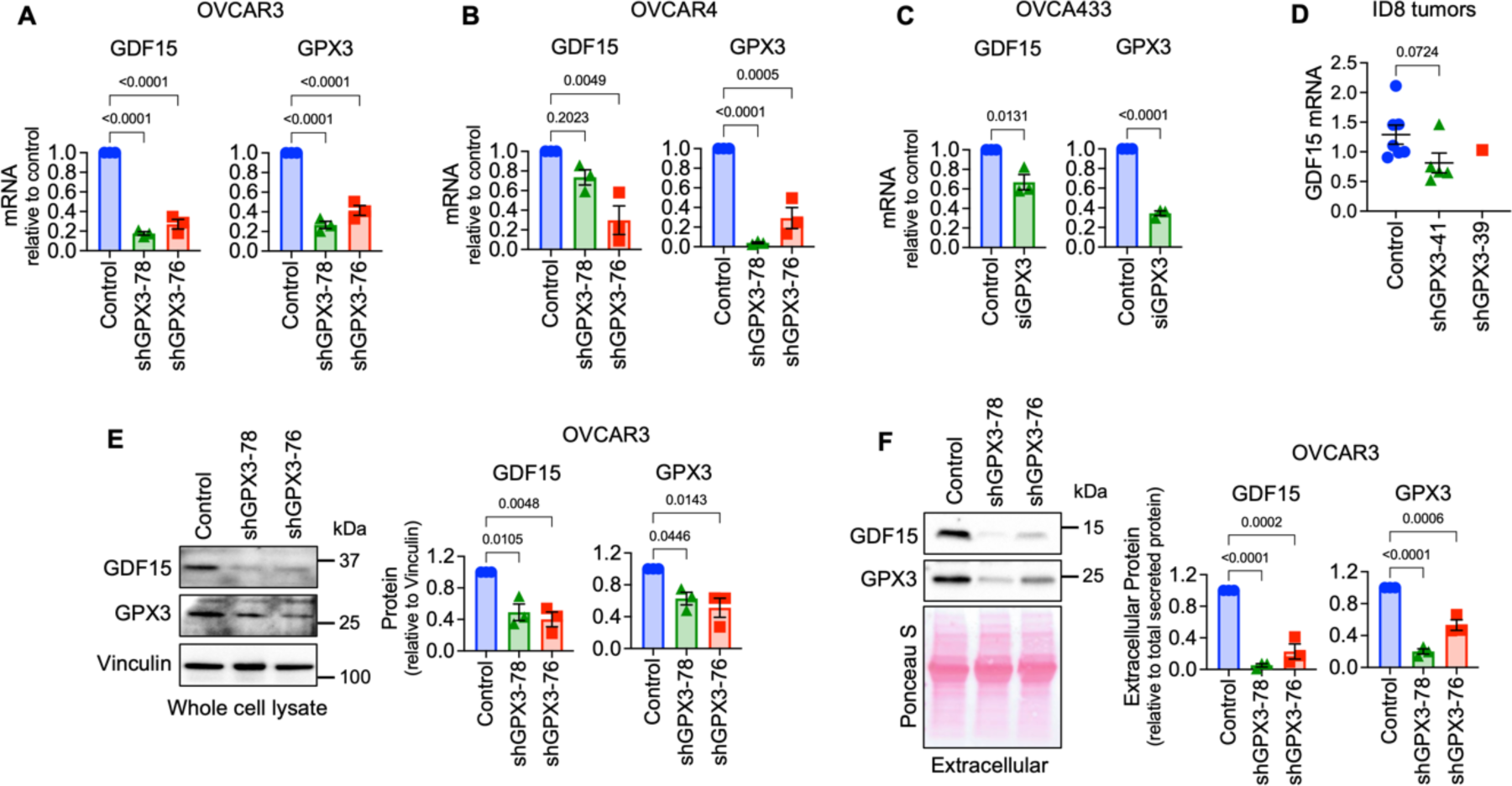
Validation of GPX3-dependent GDF15 gene expression. A. Stable shRNA mediated knock-down of GPX3 in OVCAR3 leads to decreased GDF15 mRNA expression (n=3, One Way ANOVA, Tukey’s post-test). B. shRNA-mediated knock-down of GPX3 in OVCAR4 cells results in decreased GDF15 mRNA expression (n=3, One Way ANOVA, Tukey’s post-test). C. Transient siRNA-mediated GPX3 knock-down in OVCA433 cells leads to decreased GDF15 mRNA expression (n=3, t-test). D. GDF15 mRNA expression in tumors from ID8 cells (from Figure 1), assessed by semi-quantitative real time RT-PCR (Kruskal-Wallis test). E. shRNA mediated knock-down of GPX3 in OVCAR3 cells decreases intracellular and F. Extracellular GDF15 protein expression (n=3, One Way ANOVA, Tukey’s post-test).

Of the 61 differentially expressed genes (DEGs) displaying a log 2-fold change >1 (p adjust <0.05; Figure 2E, Supplemental Data Table 1) the top protein coding gene downregulated following GPX3 knock-down was GDF15. GDF15 is a stress-responsive member of the TGF-β growth factor family, that has established roles in regulating metabolism, food intake and energy expenditure (10) (Figure 2E). Serum GDF15 levels are often elevated in cancer patients (11–14) and in a number of metabolic disorders, including non-alcoholic fatty liver disease (15), which was a top GSEA pathways identified to be dependent on GPX3 knock-down (Figure 2B).

### GDF15 is regulated by GPX3

Of relevance to ovarian cancer, GDF15 is elevated in serum from ovarian cancer patients (12), is highly expressed in ovarian tumor tissues (16), and is associated with chemoresistance (17, 18) and immune escape (19). Moreover, GDF15 induces p38, Erk1/2, Akt and NF-kB signaling in ovarian cancer cells (16, 20). Thus, we validated that GPX3 regulates *GDF15* mRNA expression by sqRT-PCR in high grade serous cell lines OVCAR3, OVCAR4, and OVCA433 using multiple sRNA hairpins and siRNA (Figures 3A-C). We were unable to statistically assess if lack of tumor burden observed in ID8 cells transduced with the shGPX3-39 hairpin is due to decreased expression of *GDF15* mRNA *in vivo*, due to the lack of sufficient amount of tumor tissue for analysis in this group (Figure 1). However, even in ID8 tumors with ∼20% knock down of GPX3 with hairpin shGPX3-41, *GDF15* mRNA expression in isolated omental tumors was lower than controls (Figure 3E). Further, GPX3 was necessary for intracellular and secreted extracellular GDF15 protein expression (Figure 3E&F). These data collectively demonstrate that GPX3 is an upstream regulator of the soluble growth factor GDF15.

### GPX3 expression is positively associated with regulatory T-cell and macrophage signatures in ovarian cancer specimens, and significantly correlates with PD-L1 expression

Cell intrinsic effects on GPX3 loss were primarily related to cellular survival in response to stress, as demonstrated by our observations that GPX3 loss inhibits single cell survival in clonogenicity assays (Figure 1B, & OVCAR3 data see (4)), and in response to exogenous oxidative stress as we previously reported (4). Using two independent analyses to survey TCGA data for immune cell expression profiles, we also found that *GPX3* mRNA expression was significantly associated with macrophages and regulatory T-cell (T-reg) and activated NK cells in ovarian cancer specimens (Figure 4A&B), suggesting that high *GPX3* expression correlates with specific tumor immune cell infiltrates. Utilizing the immune cell gene expression signatures established by Tamborero et al. (21) we found significant positive association between *GPX3* expression and macrophages, CD8+ T-cells and T-regs, and a negative association with T-helper cells (Figure 4A). Similarly, Cibersort analysis (Timer2.0 (22)) showed that T-regs, M2 macrophages and activated NK cells were positively associated with *GPX3* mRNA expression (Figure 4B). Moreover, *GPX3* expression in ovarian cancer specimens from TCGA significantly correlated with *CD274* (PD-L1) and *FOXP3* mRNA expression (Figure 4C). No correlation to *PDCD1* (PD-1) expression was found. Moreover, we could demonstrate that FOXP3 and PD-L1 expression also strongly and significantly correlated to *GPX3* mRNA expression in tumors derived from ID-8 cells (Figure 4D). These data suggest that GPX3 expression in ovarian tumors is potentially associated with “hot” immune signatures, T-reg infiltration and suppression of anti-tumor lymphocytes.

**Figure 4.**
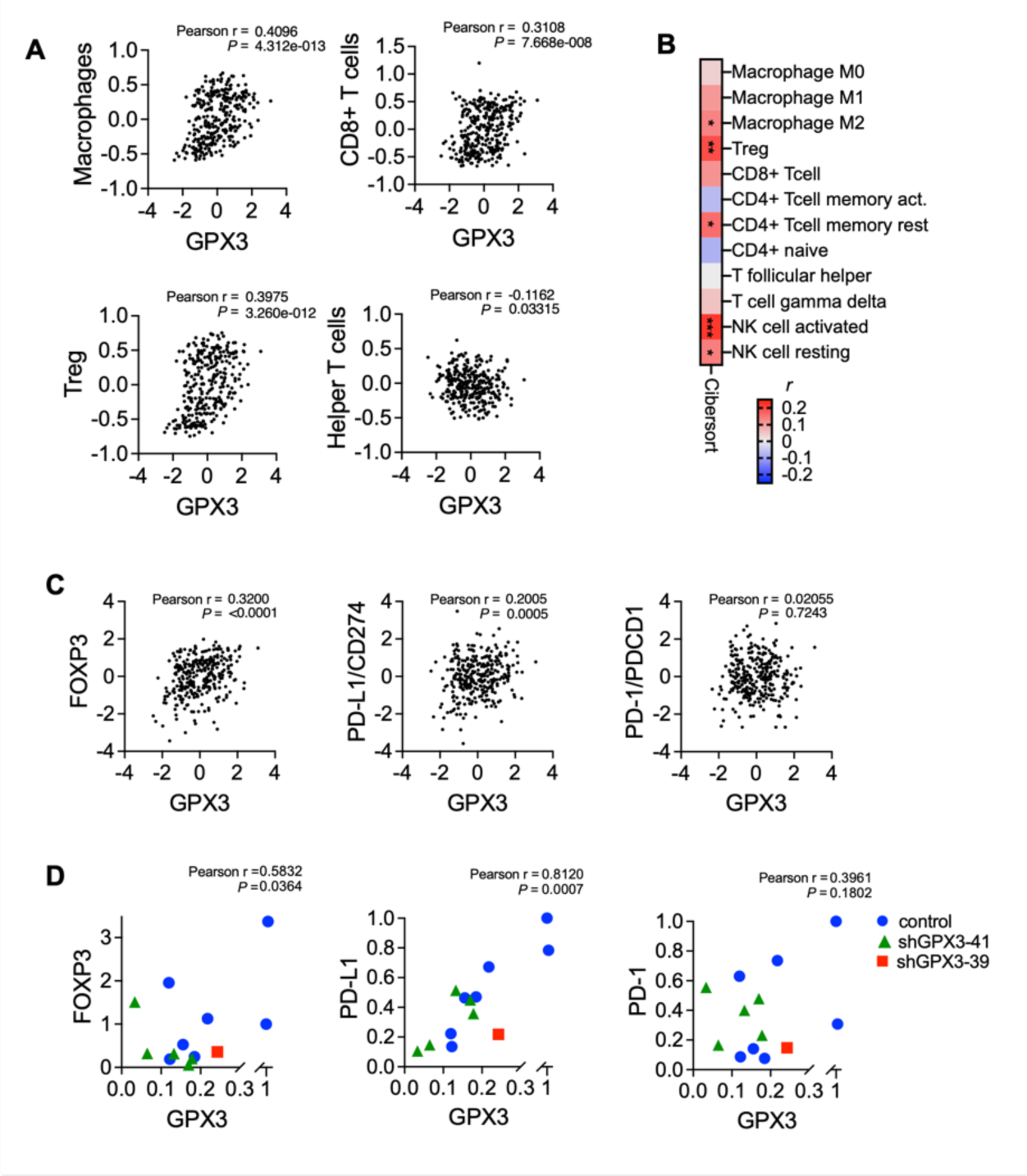
GPX3 expression is associated with immune cell infiltration and PD-L1 expression. A. Immune cell gene expression signatures established by Tamborero et al. (16) were correlated to GPX3 expression (z-score) in TCGA serous ovarian cancer specimens. GPX3 expression has significant positive correlation to macrophages, CD8+ T-cells and T-regs, and significant negative correlation with T-helper cells (Pearson r). B. Cibersort analysis of GPX3 expression to immune cell signatures in TCGA serous ovarian cancer specimens (Timer2.0, CIBERSORT-ABS, Pearson r, *P<0.05; **P<0.01; P<0.001). C. Expression of GPX3 mRNA in TCGA serous ovarian cancer specimens positively correlates with FOXP3 and PD-L1 expression (z-scores, n=300 specimens from TCGA with RNAseq data). D. Expression of GPX3 mRNA in ID8 tumors (from Figure 1) positively correlates with FOXP3 and PD-L1 expression.

## Discussion

Here we further investigated the role of tumor cell produced GPX3 in ovarian cancer, demonstrating that GPX3 is necessary for *in vivo* tumor growth in a syngeneic immunocompetent model (Figure 1). High GPX3 expression in ovarian tumor specimens, including high grade serous and clear cell subtypes has been reported by us and others, and shown to be associated with poor patient outcome (4, 8, 23, 24). Moreover, bioinformatics analyses have demonstrated that GPX3 expression correlates to specific tumor immune cell signatures. Similar to our observations that GPX3 expression is associated with CD8+ T-cell and macrophage infiltration in ovarian cancer, GPX3 expression correlates with immune cell infiltration and immune checkpoint marker expression in gastric cancer (25). Moreover, GPX3 was identified as one of 7 DEGs predictive of poor patient survival in ovarian tumors with high immune and stromal scores (7). We find that *GPX3* mRNA correlates strongly with T-reg/FoxP3 expression as well as the immune check point protein PD-L1. Interestingly, in lung adenocarcinoma, a cancer where high GPX3 expression in tumors is conversely predictive of better patient outcome, GPX3 expression was shown to be significantly correlated to PD-L1 expression and identified as a predictive marker of immune check point inhibitor (ICI) response (26). EREG was also identified as a candidate ICI predictive marker as it strongly correlated to PD-L1 expression. Interestingly, we also found EREG to be under the regulation of GPX3 and correlated to GPX3 expression in HGSOC TCGA specimens (Figure 2&3).

Although we focused on manipulating GPX3 expression in tumor cells, it is possible that GPX3 in the extracellular tumor environment (TME) is also derived from tumor associated cells. As such, it was recently shown that alveolar type 2 epithelial cells near the tumor interface contribute to GPX3 expression in the TME and that this is necessary for metastatic colonization of the lung, by inhibiting CD4^+^ T cells and supporting T-reg cell proliferation, suggesting that high GPX3 in the TME, contributed by tumor associated cells is also immuno-suppressive (27). This correlates with our observations that high GPX3 expression in HGSOC correlates with enhanced PD-L1 expression and T-reg (FoxP3) infiltration signatures. In recent work it was also found that GPX3 is strongly expressed in stromal enriched colorectal cancer where its expression promotes chemoresistance (28). Querying publicly available single cell RNAseq data (TISCH2.0) we find that GPX3 in HGSOC tumor samples is primarily expressed by metastatic cells and myofibroblasts. How tumor associated cells contribute to the extracellular GPX3 milieu and its consequence on the TME in ovarian cancer remain to be determined.

Our previous study demonstrated that tumor cell produced GPX3 expression enhanced OVCAR3 clonogenicity and anchorage-independent survival. We found that GPX3 provides survival advantages to tumor cells when exposed to exogenous oxidative stress, either through the addition of high dose ascorbate, which reacts with iron to produce H_2_O_2_, or in response to culture in patient derived ascites fluid, a tumor medium with potential high source of iron that correlates with oxidative stress (4). In addition, we now demonstrate that GPX3 is necessary for maintaining a transcriptional program associated with pro-tumorigenic signaling pathways (Figure 2). We find that the expression of GDF15, a mitokine that has pro-proliferative and pro-metastatic growth factor functions in cancer, is strongly GPX3-dependent (Figure 3). Numerous studies have shown that GDF15 activates signaling pathways, including Smad2/3, Erk and Akt signaling in tumor cells (14). Interestingly, these signaling pathways were also significantly decreased following GPX3 knock-down in OVCAR3 cells. Future studies will determine if GDF15 is the main factor downstream of GPX3 driving these pro-tumorigenic signaling changes. Although GDF15 belongs to the TGF-β family, it is not thought to directly signal through TGF-β receptor (10). Tumor produced GDF15 is also linked to cachexia via its activation of the brain localized receptor GFRAL (GDNF Family Receptor Alpha Like) and interaction with RET to induce ERK and Akt signaling, which initiates appetite suppression (29). Since GFRAL expression is limited to the brain, it remains unclear which receptor is the primary target of GDF15 in tumor cells. Our RNAseq data demonstrate that many metabolic pathways are decreased following GPX3 knock-down, in particular fatty acid, amino acid and mitochondrial respiration, and it is possible that this is also linked to the metabolic function of GDF15, which requires further investigation.

Due to its increased expression in many tumors and serum of cancer patients, GDF15 has been studied as a potential tumor biomarker, including ovarian cancer (12, 16). Interestingly, besides its intrinsic pro-tumorigenic role, which has been linked to induction of p38, Erk1/2, Akt and NF-kB signaling in ovarian cancer cells (16, 20), GDF15 has also been implicated with the regulation of inflammation and tumor immune escape (14). GDF15 was shown to inhibit the function of Dendritic cells (DC), leading to decreased DC dependent T-cell stimulation and lack of cytotoxic T-cell activation in a liver cancer model (30). In ovarian cancer, GDF15 is associated with higher DC infiltration, but lack of activated DCs (19). Moreover, GDF15 expression correlates positively with PD-1+ T-cells in plasma of lung cancer patients (31), and enhances T-cell exhaustion and increases T-reg cell expansion via CD48 mediated FoxP3 stabilization in a hepatocellular carcinoma model (32). In gall bladder cancer, GDF15 has been shown to increase PD-L1 expression in a PI3K/Akt/Erk dependent manner to result in immune suppression (33). In addition, a recent study has shown that GDF15 suppresses responses to anti-PD1 immune check point therapy, as GDF-15 impairs T-cell endothelial adhesion and thus recruitment to the tumor, with T-reg recruitment being less affected (34). Thus, GPX3 and GDF15 appear to share similarities in their correlation to tumor immune cell signatures. Although we show that GPX3 is necessary for the expression of GDF15 by tumor cells it remains to be determined if GPX3 is the necessary upstream regulator of GDF15 in the context of the tumor infiltrating lymphocytes.

Another interesting parallel between the actions of GDF15 and GPX3 are the potential role on platinum resistance. GDF15 levels are increased post-chemotherapy in ovarian cancer patient effusions (35), high expression of GDF15 serum levels can predict patient response to platinum based therapies (18), and GDF15 knock-down can increase chemosensitivity in chemoresistant cells (17). Similarly, GPX3 has been shown to confer resistance to cisplatin in colorectal cancer (36) and ovarian clear cell carcinomas (8), although high GPX3 expression in ovarian cancer effusions was not significantly associated with chemotherapy response (37).

GPX3, like many antioxidant enzymes, has dichotomous functions in cancer, able to act as both a tumor suppressor during oncogenesis, and a tumor promoter during tumor progression. This dichotomous function is linked to its role as an antioxidant enzyme in protecting both normal cells during oncogenesis and tumor cells during tumor progression from oxidative stress. It is likely that part of the immune-modulatory function of GPX3 is associated with its antioxidant activity in the TME, making it a more permissive environment for T-reg expansion, for example. A limitation to our work is that further studies are required to elucidate the mechanisms by which GPX3 regulates GDF15 expression. GDF15 expression is often induced in pathological conditions including inflammatory diseases, during cardiac and renal failure and in cancer patients (10), and has been shown to be induced in response to stress, including hypoxia, mitochondrial dysfunction (38), engagement of the integrated stress response and pro-inflammatory cytokines (39) and PPARγ agonists (40). Moreover, it was initially identified as NSAID activated gene −1 (NAG-1) due to its inducibility by NSAIDS (41). Thus, it was somewhat surprising that loss of the antioxidant enzyme GPX3, and presumably an increase in resultant cellular stress, decreased GDF15 expression. However, pathway analysis of RNAseq data following GPX3 knock-down strongly correlate with regulatory pathways that have been implicated in GDF15 expression including unfolded protein response, PERK and ATF4/6 pathways (15). Interestingly, in the present work, we did not challenge cells with exogenous sources of oxidative stress, nor did we observe significant enrichments in pathways associated with oxidative stress following GPX3 knock-down in our RNA sequencing analysis (Figure 2). It is thus possible that GPX3 has alternate functions besides scavenging damaging peroxide species such as H_2_O_2_ in response to extracellular oxidative stress. One potential alternative is that GPX3 acts to alter signaling lipid peroxide species in the TME that could contribute to the transcriptional changes downstream of GPX3. As such it is possible that GPX3 plays a role in the synthesis pathway of hydroxyeicosatetraenoic acid (HETEs) and oxoeicosatetraenoic acid (oxo-ETEs) derived from arachidonic acid, as the peroxidase responsible for the reduction of HpETE to HETE, which is subsequently converted to oxo-ETE (42). HETEs and oxo-ETEs are pro-proliferative in cancer cells and both are able to activate PPARγ (42–44), a transcription factor which is able to activate GDF15 expression (40).

In conclusion, we find that GPX3 is necessary for optimal *in vivo* tumor growth in a syngeneic ovarian cancer model and that expression of the pro-tumorigenic and immune modulatory growth factor GDF15 is dependent on GPX3 expression. The strong association of GPX3 expression with the immune checkpoint marker PD-L1, suggests that GPX3 expression could be predictive of immune check point inhibition response. Moreover, given that GPX3 and GDF15 are both extracellular proteins, their targeting in the TME could be a novel therapeutic opportunity.

## Acknowledgements

The authors would like to thank Sara Shimko, Weihua Pan and Benjamin Yankaski for technical assistance. This work was supported by the U.S. National Institutes of Health grants R01CA242021 (N.H.) and R01CA230628 (N.H. & K.M.). Sierra White is supported by training grant T32HL110849.

## Author contributions

C.C., and Y.Y.C. contributed to conceptual and experimental design, carried out experiments and data analysis, prepared figures and wrote the manuscript. S.K., S.W., P.W.T., A.T.E., Z.J. assisted with experiments and manuscript editing. K.M.A., K.M. and R.P. contributed to conceptual and experimental design, data interpretation and manuscript editing. N.H. supervised and conceived the study, contributed to experimental design, assisted in data analysis, prepared figures, wrote and edited the manuscript.

## Supplemental Figures

GPX3 supports ovarian cancer tumor progression *in vivo* and promotes expression of GDF15. Chang *et al*., 2024

**Supplemental Figure 1.**
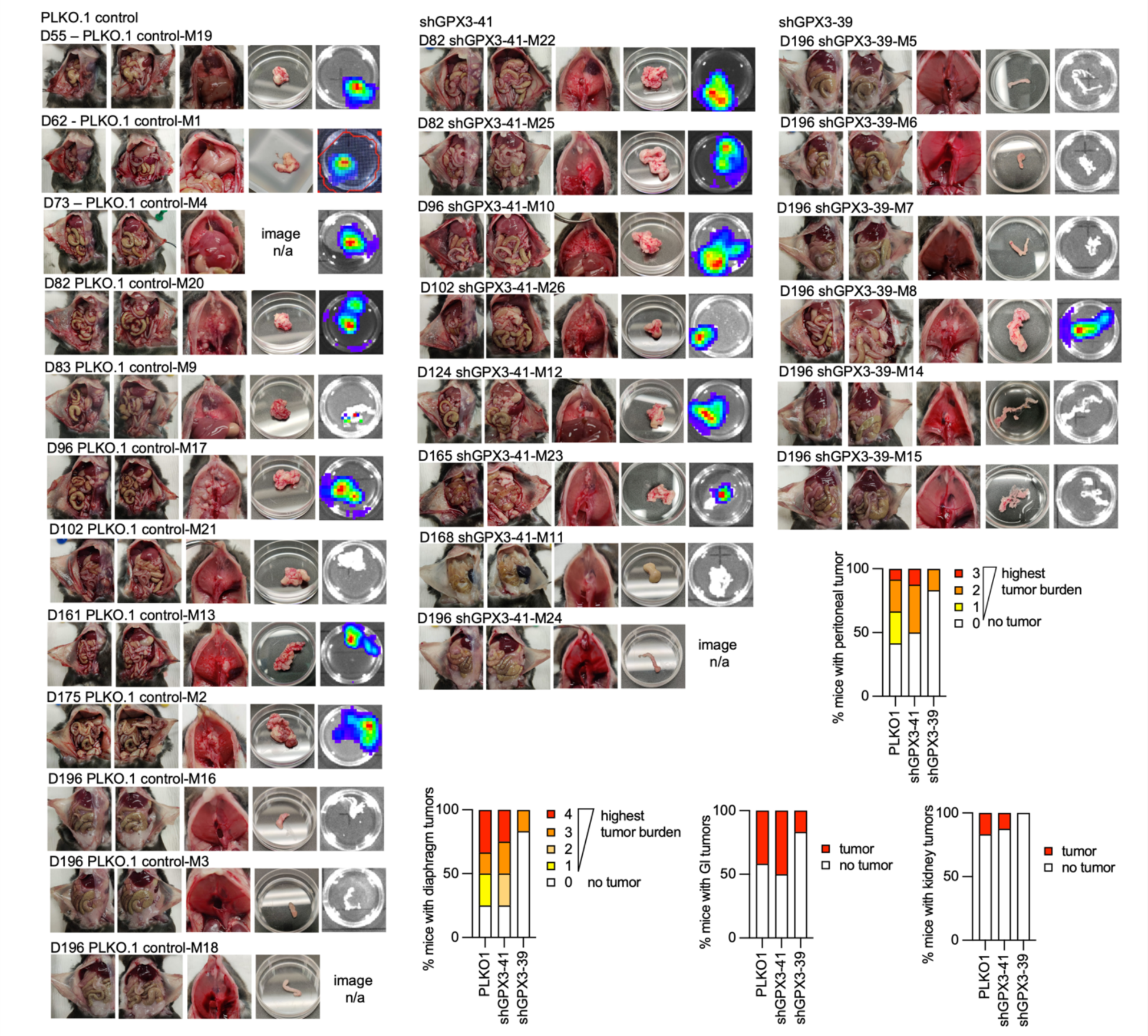
Effects of GPX3 knock-down on IP tumor burden in a syngeneic mouse ovarian cancer tumor model. ID8 tumor cells transduced with pLK0.1 control, shGPX3-41 or shGPX3-39 shRNAs targeting GPX3 and cells injected IP into female C57BL/6J mice. GPX3 knock-down by shRNA-39 results in decreased peritoneal, diaphragm, GI and kidney tumors compared to control and shGPX3-41 transfected cells. Graphs show percentage of mice with indicated tumors (pLK0.1 control n=12, shGPX3-41 n= 8, shGPX3-39 n=6).

**Supplemental Figure 2.**
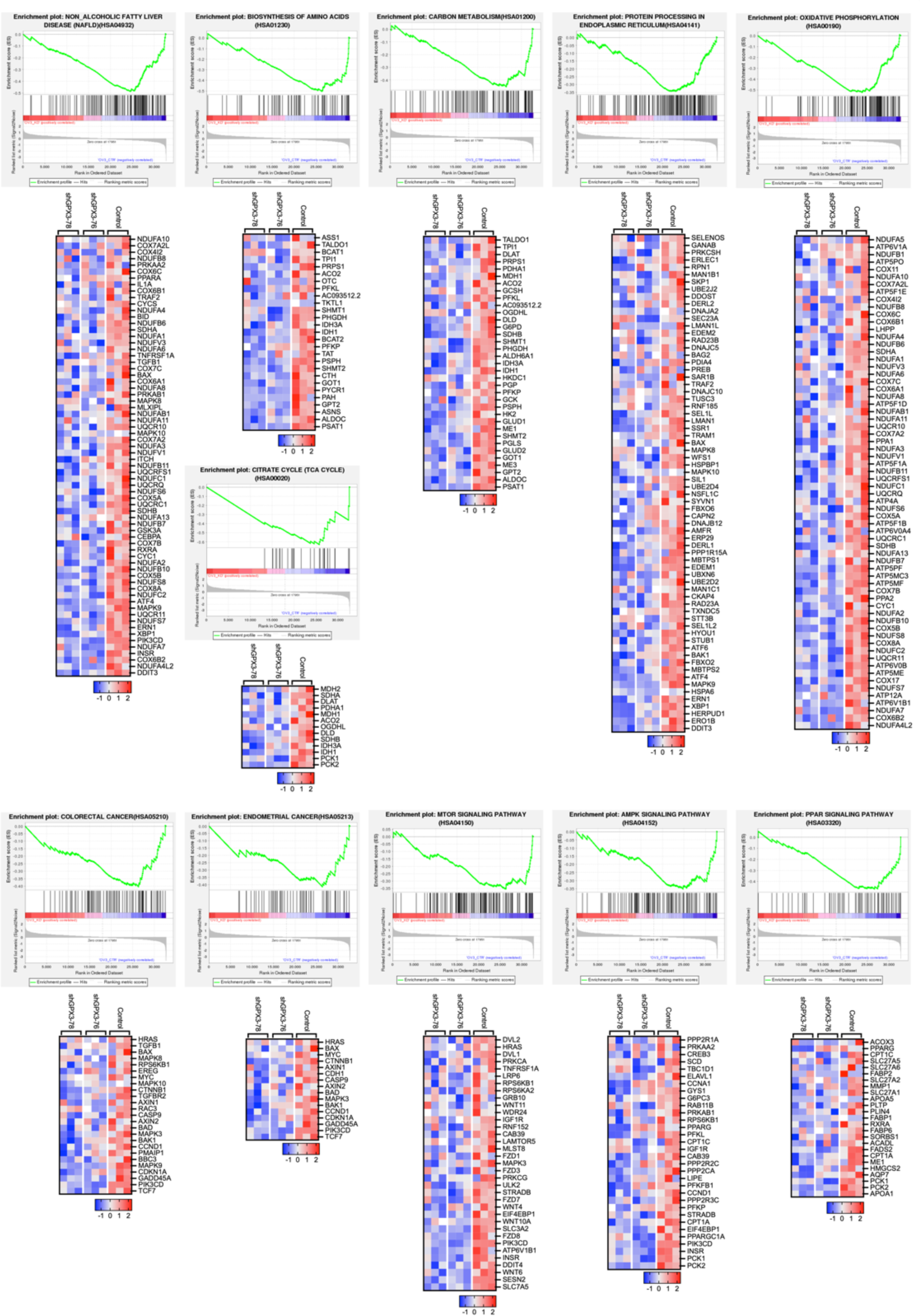
Enrichment plots of gene expression signatures downregulated following GPX3 knock-down. OVCAR3 cells were stably transduced with two independent shRNAs against GPX3 (shGPX3-78 & −76) or scramble control. GSEA was carried out on differentially expressed genes identified by RNA sequencing following GPX3 knock-down. NES are shown in Figure 2B of the manuscript. Heatmaps of core leading-edge genes of each pathway are shown (z-scores).

